## Supplementary material for "zIncubascope: long-term quantitative imaging of multi-cellular assemblies inside an incubator"

Anirban Jana,[1,2,3] Naveen Mekhlieri,[1,2] Adeline Boyreau,[1,2] Aymerick Bazin,[1,2] Nadège Pujol,[3] Kevin Alessandri,[3] Gaëlle Recher,[1,2] Pierre Nassoy,[1,2] and Amaury Badon[1,2]
[1)] *LP2N, Laboratoire Photonique Numérique et Nanosciences, Univ. Bordeaux, F-33400 Talence, France*
[2)] *Institut d' Optique Graduate School & CNRS UMR 5298, F-33400 Talence, France*
[3)] *Treefrog Therapeutics, F-33600, Pessac, France*

(*Electronic mail:)

(Dated: 5 March 2024)

### I. BIOLOGICAL SAMPLE PREPARATION

#### A. Human induced pluripotent stem cells (hiPSCs)

hiPS cells purchased from Corriell Institute (AICS-0023) are cultured using standard protocols in T25 plastic flasks (Corning-Ref: 353109), coated with Matrigel (Corning-Ref: 354234). Once at a confluency of 75-80 %, the cells are detached using Acutase (Stemcells-Ref: 07920) and resuspended in culture media at the required concentrations. The culture medium used for cell, spherical and tubular capsules culture is mTeSR™ Plus (Stemcells-Ref: 100-0276). The detached cells are encapsulated with Matrigel (Corning-Ref: 354234) in equal volumes of cell solution and the Matrigel. In this case of capsules and tubes, the alginate assemblies of hiPSCs are rinsed with DMEM (Biowest L0102) to remove excess calcium and cultured in mTeSR™ Plus culture medium. ROCK inhibitor Y-27632 dihydrochloride (Tocris-Ref: 1254/10) at 10 mM is added and replenished at 1/1000 dilution every day for the first 48 hours after encapsulation. After 48 hours, culture media with Penicillin-streptomycin (Gibco™ 15140122) in 1:100 dilution is used and replaced every 48 hours.

#### B. Yeast cells

For encapsulation, the yeasts *Saccharomyces cerevisiae* (Fermentis-Ref: SafAle™ S-04) are suspended at a concentration of 10 mg/ml in a solution of 300 mM sorbitol (Applichem 548653) before encapsulation. Once encapsulated, the capsules are cultured in Yeast Peptone Dextrose (YPD) broth (Sigma-Ref: Y1375). The media is unchanged during the imaging time period.

#### C. Spherical and tubular cellular capsules

We used the previously developed Cellular Capsule technology[1,2] to encapsulate hiPS and yeast cells either in spherical or in tubular hollow hydrogel capsules[3]. In brief, the spherical capsules are made using a co-extrusion microfluidic device that generates a jet with a core-shell geometry. Owing to the Rayleigh-Plateau instability, the jet breaks up into droplets with a shell of sodium alginate solution and a core of cells suspended in an extra-cellular matrix[4]. These fall into a 100 mM $CaCl_2$ and instantaneously crosslink to form a hydrogel. We thus obtain spherical shells (i.e. hollow spheres) with cells encapsulated in the aqueous core and the shell serving as a protective membrane against manipulation but permeable to nutrients. The size of the capsules is mainly set by the diameter of the nozzle of the microfluidic coextrusion device. In the experiments reported here, the capsules has an inner diameter of 250±50 µm and an outer diameter of 400±100 µm. The tubular capsules are made using the same co-extrusion device; however, the composite liquid jet is prevented from undergoing the Rayleigh-Plateau instability by dipping the device into the calcium bath to suppress significant surface energy mismatch. The tubes could be as long as meters with an outer diameter of 250 µm. As for spherical capsules, the cellular concentration inside the core of the tube can be tuned for different experiments. The tubes are maintained in culture using the same protocols as in the case of capsules.

One peculiarity of the technique that we exploited here is the fact that alginate gels stick to positively charged surfaces. We thus used petri dishes coated the poly-lysine (Gibco-Ref: 16021412) before seeding the capsules to avoid undesired motions of the samples during acquisition, especially upon culture medium exchange.

### II. IMAGE ANALYSIS

#### A. Capsules

The images obtained from the acquisitions are processed using ImageJ (FIJI[5]) and quantitative analysis is performed in Matlab R2023 (MathWorks). The .m files are provided as digital supplementary material. To analyze the growth of a cyst, from a z-stack, a suitable focal plane is chosen for analysis. This focal plane varies for different cysts. The whole pipeline is illustrated in figure S1. Concretely, the full-field images are cropped into regions of interests (ROIs) to be analyzed individually. In the case of spherical capsules, a binary mask of the cyst is constructed in FIJI using simple threshold and binary operations of `erosion`, `dilation`, and `fill holes`. A local rectangular (ROI) is defined using the centroid of this mask as the lower left corner of the rectangle. The width (w in Fig. S1(a) ) of this local ROI is defined using the half-width of

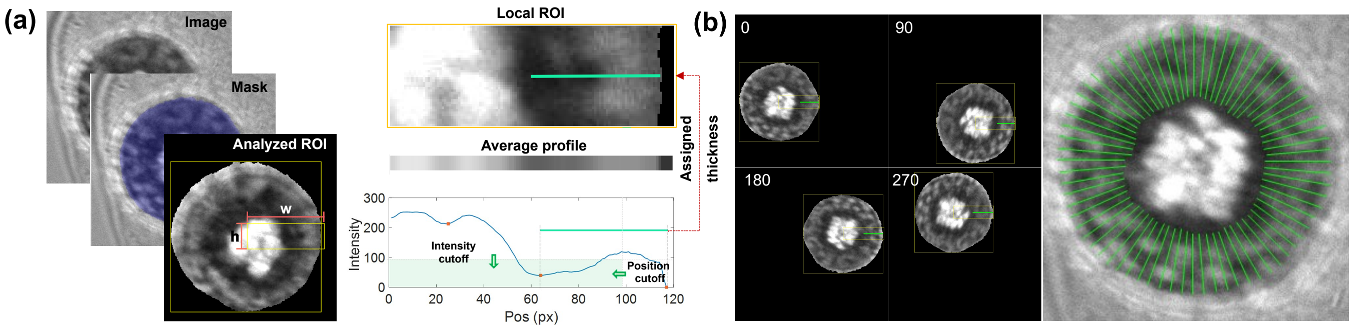


Fig. S1. Detection of cyst thickness. (a) Image analysis pipeline to extract the thickness of the cyst in the local ROI using preset thresholds on the local minimas of the averaged intensity profile. (b) Four angular positions of the mask and the detected thickness of the cell (layer (in green in each snippet). Detection of the thickness of the cell layer is performed by rotating this mask around 360°.

the bounding rectangle of the global mask with 5 pixels added to this value. The height (h in Fig. S1(a)) of this ROI is preset at 20 pixels to obtain a locally averaged radial intensity profile to prevent detection of smaller dark artefacts in the cell layer (refer to Fig. S1).

Local quantification of the thickness of the cyst is performed on this locally masked region of the gray-scale image. Since the cyst has a central cavity (lumen), the edge of the cell layer appears as a dark band that is manifested as a local minimum in the average radial intensity profile of this ROI. The local minima in this profile are identified with a preset prominence value (Matlab function: `islocalmin (A, P)`). A threshold is defined in both intensity values and position on this averaged profile to extract the edge of the cell layer facing the lumen. The threshold in intensity depends on the global intensity values, however the same value is used for a single FOV since illumination is homogeneous in all the cases. The cut-off in position is set at a fixed distance from the outer edge of the cyst, implying that the distance of the lumen edge from the centroid is below this cut-off distance. The outer edge of the cyst in the ROI is the first non zero pixel value position from the right extremity of this profile curve. These edge positions are assigned to the central pixel line of the cyst of the local ROI. Using the centroid of the mask and these two positions on the cell layer, we quantify the lumen radius and the thickness of the cell layer in this local ROI. To obtain the thickness of the cell layer around the cyst, the gray-scale image and the mask is rotated. The local ROI is calculated at each rotation step and the above-mentioned operations are carried out. The rotation step can be smallest positive normalized floating-point number in MATLAB, however for computational costs, it has been set at a step value of 5. This method ensures that the non-uniformity in the lumen radius and the cell layer are well captured in the calculations.

### B. Tubes

A similar analysis strategy is utilized to estimate the thickness of hiPSCs tubes (section III). A mask of the cyst is created using binary operations at each time step and a local rectangular ROI is extracted to analyze the thickness. In the case of hiPSCs tubes, the ROI is translated along the axis of the tube to detect the radial thickness of the tubular cyst. Similarly, to detect the thickness of the cyst at the free tip (axial position), the ROI is translated in the radial direction (Fig. S2(a)). The thickness is detected every 2.5 µm along the radial and axial directions. The thickness in the radial position as shown in the analysis Fig. S2(b) is always calculated in a region located at a fixed distance from the free tip of the capsule. The ROI is defined using the central axis of the tube as a reference with a preset width and height to ensure proper detection of the edges of the cyst and the lumen. The edge of the lumen is detected as the local minima in the local ROI using thresholding techniques, as in the case of spherical capsules, on the mean intensity profile of the ROI. The outer edge of the cyst in the ROI is the first non zero pixel value position from the outer-extremity of this profile curve.

### C. Yeast

In the case of yeast cells *Saccharomyces cerevisiae* growing in capsules, it is observed that they proliferate as aggregates, hence we did not investigate the structural details at the scale of the cellular assemblies. Owing to a high homogeneity in the illumination of the full field of view, a global mask is developed using binary thresholding of the intensity. Individual capsules with their masks are cropped for analysis from different regions of the field of view. The isolated masks are dilated to circumscribe the regions of interest and properties of this mask are analyzed. Since the aggregates are non-spherical and in-homogeneous, a volume calculation using the length scale of the 2D projected area is inaccurate. Hence, an estimation of the depth of the aggregate is necessary to obtain the volume. Following Beer-Lambert's law, the depth of each pixel is calculated using the intensity value of each pixel with the background intensity value (in the region without capsules, $I_0$=140) following the equation in Supp Fig. S3(d). The absorbance constant of the Beer-Lambert equation is assumed to be identical for each pixel and constant w.r.t time. The summation of the volume of each pixel yields the volume of the

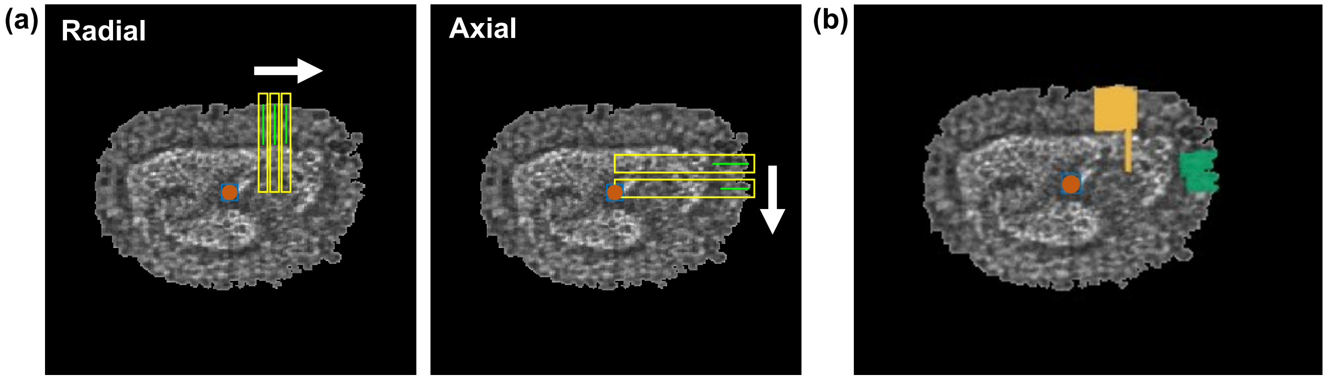


Fig. S2. Detection of thickness of tubular cysts. (a) Translation of the local ROI to trace the thickness in the radial position (here shown at steps of 10 µm for clarity). (b) The thickness detected at the radial and axial positions using the translating ROIs at steps of 2.5 µm at t=0 for the data shown in section III B, Fig. 7.

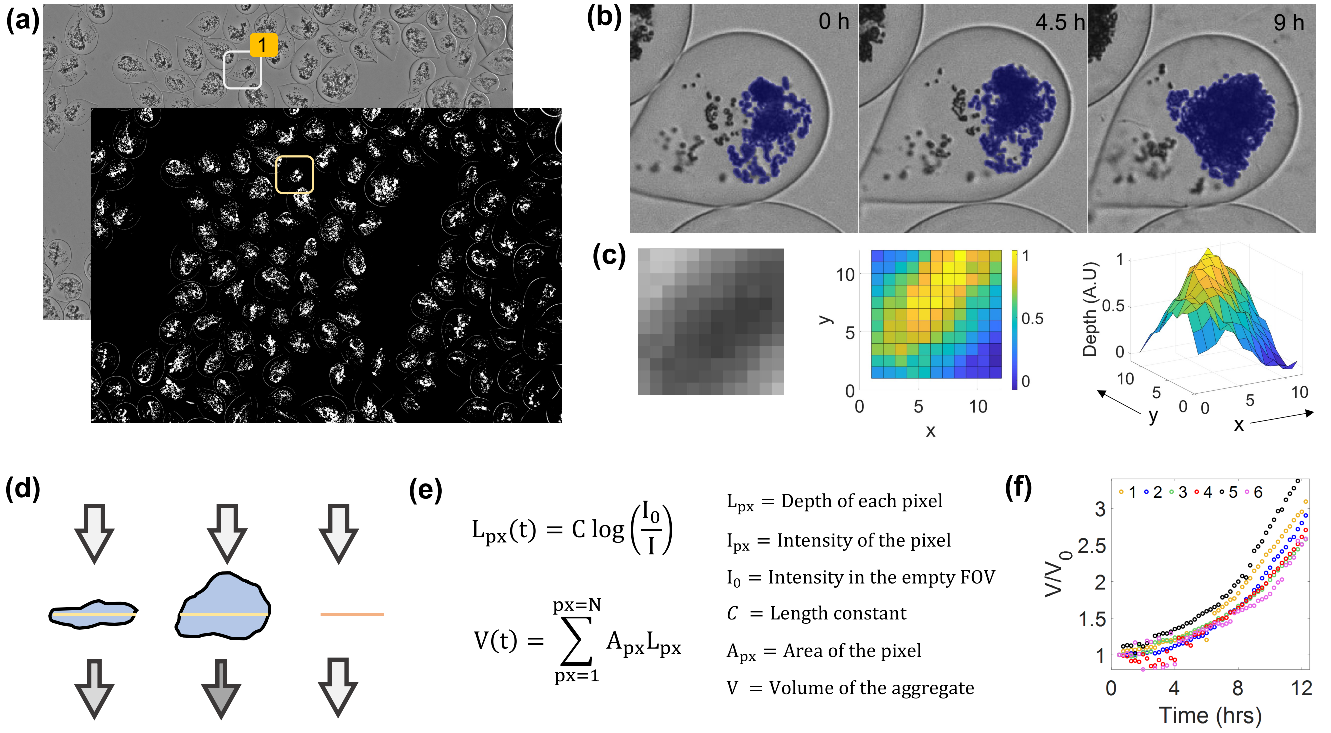


Fig. S3. Detection of volume of yeast aggregates. (a) Image analysis pipeline to develop the global mask of the FOV to analyze growth of yeast aggregates. (b) Detection of the area occupied by the yeast cells at different time steps. (c) Depth calculation for each pixel using Beer-Lambert's law. The color map represents a normalized depth distribution of a small region of the mask at a given time. (d) Schematic of how depth of an aggregate with identical projected area affects the mean intensity of the light of the ROI. (e) Estimation of the volume of the aggregate using Beer-Lambert's law. (f) Variation of the estimated volume (normalized with the volume at t=0) with time for the aggregates shown in Fig. 9 (a)).

aggregate at a given time (Supp. Fig. S3(c)). Normalization of the volume of the aggregate with the value at t=0 ensures that the constants from the equation can be removed from the final data.

### III. Setup to Measure Accuracy

To measure the performance of the motor driven z-stack capabilities of the zIncubascope, we have designed a setup to quantify it. Using an LED and collimator lens (LED-L in Fig. S4), a stage (STAGE in Fig. S4) is illuminated as shown.

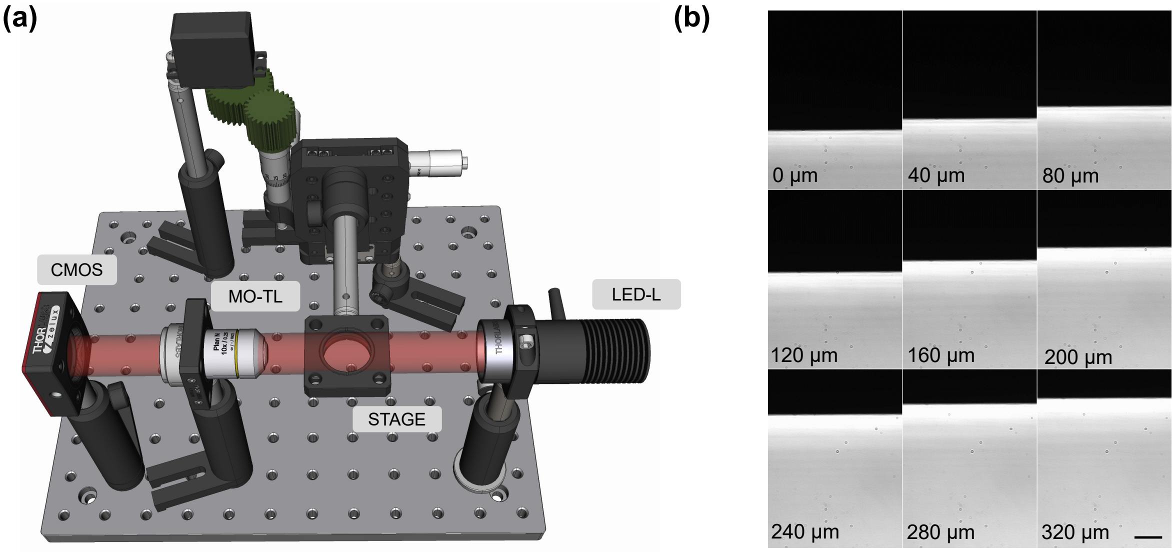


Fig. S4. 3D rendering of the optical setup. (a) CAD scheme of the setup used to measure the step displacement of the stage for controlled motions of the motor. (b) Truncated timelapse of an automated run performed with steps of 5 degrees of rotation of the motor (10 µm displacement step). Note: The lower edge of the stage is imaged, and the position of the lower edge is monitored as it is displaced upwards.

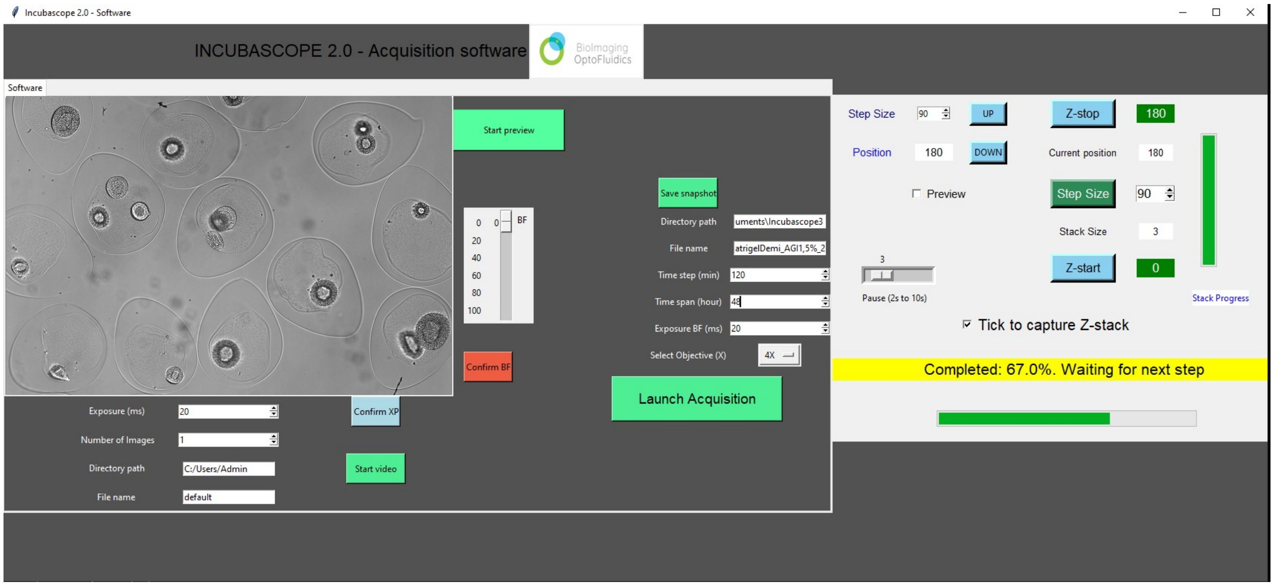


Fig. S5. Python Interface to control the functionalities of the zIncubascope.

This light is collected using a 10X objective (Zeiss-Ref: 13-816 10X ZEISS A-Plan Objective) and tube lens on a CMOS (Thorlabs-Ref: CS165MU1/M). The camera has a pixel size of 3.45 µm and using a tube lens of 125 mm combined with the focal length of the objective (16.5 mm) provides a resolution of 0.45 µm per pixel. Since the shortest displacement expected is 10 µm, this resolution is sufficient to measure the movements with high accuracy.

### IV. ACQUISITION CODE

All experiments were performed using a graphical user interface developed in Python available in the BiOf GitHub repository (github.com/BiOflab/zIncubascope). We designed an ergonomic interface (Supp. Fig. S5) that provides a preview window, control of the LED power, camera acquisition parameters, motor control and allows to enter the parameters for a time lapse (time step, duration). To further reduce the cost as well as the physical size of the system outside the in-

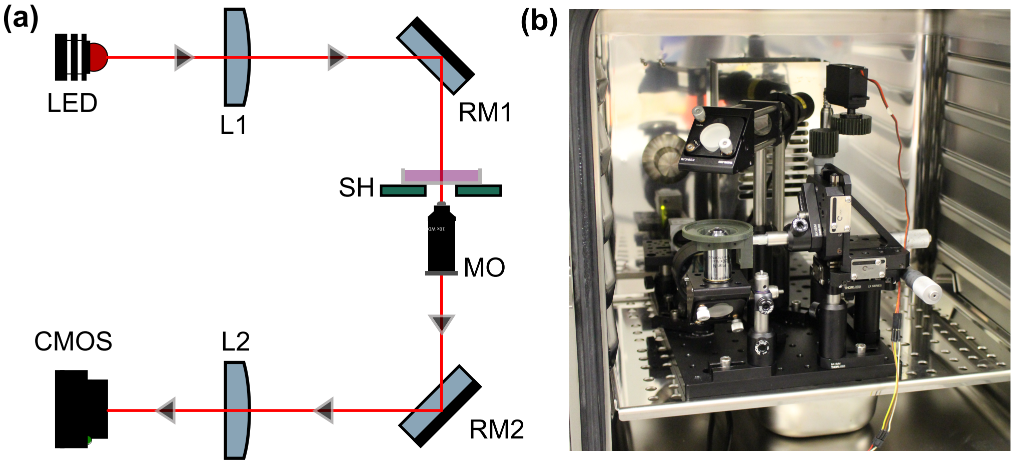


Fig. S6. Experimental setup. (a) Illumination pathway of the zIncubascope. Refer to Fig. 1 for more details. (b) A picture of the setup in a tabletop incubator.

cubator, we also developed a version of the acquisition code compatible with Raspberry Pi 4 systems. For more details refer to our GitHub page[6].

### V. EXPERIMENTAL SETUP

On figure S6, we provide a 2D sketch of the experimantal apparatus as well as a picture of the zIncubascope inside a 118L incubator.

### VI. VIDEOS

- Supp Video 1: Timelapse of an hiPSCs cyst growing in a spherical capsule (cropped from the full FOV) from Day 1 to Day 3. (section III A)
- Supp Video 2: Timelapse of hiPSCs cysts growing in spherical capsules from Day 4 to Day 6. (section III A)
- Supp Video 3: Timelapse of 3 hiPSCs cysts growing in a spherical capsule imaged in different planes from Day 4 to Day 6. (section III A)
- Supp Video 4: Timelapse of hiPSCs proliferating in a tube of alginate from Day 1 to Day 5. (section III B)
- Supp Video 5: Timelapse of yeast proliferating in spherical capsules over 12 hours. Scale bar=500 µm. (section III C)

[6] `https://github.com/BiOflab/zIncubascope`.
